# Accurate peptide fragmentation predictions allow data driven approaches to replace and improve upon proteomics search engine scoring functions

**DOI:** 10.1101/428805

**Authors:** Ana S. C. Silva, Lennart Martens, Sven Degroeve

## Abstract

The use of post-processing tools to maximize the information gained from a proteomics search engine is widely accepted and used by the community, with the most notable example being Percolator - a semi-supervised machine learning model which learns a new scoring function for a given dataset. The usage of such tools is however bound to the search engine’s scoring scheme, which doesn’t always make full use of the intensity information present in a spectrum. We aim to show how this tool can be applied in such a way that maximizes the use of spectrum intensity information by leveraging another machine learning-based tool, MS2PIP. MS2PIP predicts fragment ion peak intensities. We show how comparing these intensities to annotated experimental spectra by calculating direct similarity metrics provides enough information for a tool such as Percolator to accurately separate two classes of PSMs. This approach allows using more information out of the data (compared to simpler intensity based metrics, like peak counting or explained intensities summing) while maintaining control of statistics such as the false discovery rate.

## Introduction

Proteomics is a field that relies heavily on mass spectrometry for the identification of proteins in a sample^1^. The identification process of the acquired fragmentation mass spectra is carried out using bioinformatics tools called search engines that match experimentally obtained mass spectra to peptide sequences ^23^.

Sequence database search engines, who are by far the most popular kind of search engine, assign sequences to spectra by generating theoretical spectra for each potential sequence, matching it to the experimental spectra and attributing a score to each match^23^. These theoretical spectra are typically very simple: for example, SEQUEST^10^ creates these by assigning an arbitrary magnitude of 50 for b-and y-ion fragment peaks, of 25 for ions with m/z equal to +-1u from the b- and y-ion fragments, and of 10 for ions associated with neutral losses of water or ammonia. These different intensities aim to represent the relative abundances of the different ion types. Even though some distinction is thus hinted at, this approach is far from representing the real differences in intensity seen experimentally.

A score is then calculated based on the match between such a theoretical spectrum and an experimental spectrum. This peptide-to-spectrum match (PSM) score is intended to allow the discrimination of truthful matches from random matches. This is done through scoring functions that can be based on the number of matched peaks, or on the sum of matched experimental peak intensity. Today, these scoring functions are typically extended with a statistical model that attempts to model the probability of obtaining a match at this score by chance^25^. The output of a search engine consists of PSMs with scores to indicate their reliability against a random match model, which typically take the form of an e-value or similar statistical metric.

While this approach delivers reliability metrics for individual PSMs, it does not address the overall number of incorrect PSMs that can be expected to occur when many PSMs are reported. To address this issue, a correction for multiple testing needs to be performed, and this is typically implemented as a false discovery rate (FDR) control that is calculated across the top-scoring PSMs for each spectrum^3,17^.

In proteomics, FDR control is not based on statistical models, but is derived from empirical data. This is achieved by searching a compound target-decoy database, in which the target database contains the protein sequences of biological interest, while the decoy database contains nonsensical sequences that are designed to accurately represent false positive identification results^19^.

In order to improve the amount of spectra identified at a fixed FDR, researchers frequently couple search engines to specially developed post-processing tools. These tools employ machine learning algorithms to separate true from false PSMs by exploiting all the information available of the PSM in the form of feature vectors. This information includes PSM scores computed by the search engine, delta scores, peptide mass, charge state, precursor mass error, fragmentation mass errors, etc. Post-processing tools have been available from as early as 2002 ^12^, and have since gained increasing popularity in the normal proteomics bioinformatics workflow.

The earliest attempts at tackling the PSM identification problem from a data-driven perspective framed it as a binary classification problem, where the positive class is represented by confident target PSMs, while the negative class is represented by decoy PSMs. One such attempt was Anderson et al.’s^2^, who trained a support vector machine (SVM) on PSMs obtained from SEQUEST^10^. The SVM was trained on thirteen PSM features (of which nine were obtained directly from the search engine’s score calculations), with the positive class of PSMs selected by expert criteria, while the the negative class corresponding to PSMs associated with incorrect proteins.

Percolator^11^ perfects this approach. While similar in principle, it introduces some crucial differences that have made it into the most popular post-processing tool in the proteomics community [The 2016]. The first of these differences was the re-framing of the problem as a semi-supervised problem: after a search engine run, the results include three rather than two classes: “negative”, represented by the decoy hits; “positive”, represented by the target hits above the FDR threshold; and “unknown”, represented by the target hits which fall under the FDR threshold. An SVM is trained on the positive and negative classes in order to learn a (scoring) function that best separates these two classes. The second crucial difference is that this new scoring function is not meant to generalize to other data sets, but only to rescore the PSMs in the current data set – including the PSMs in the “unknown” class. In this way, members of this class can be re-assigned to the positive class, and more PSMs can be recovered at the same FDR cut-off.

Here, we replace the search engine features with spectral comparison features, with the goal of overcoming post-processing tools’ dependency on the search engine’s score calculations and instead maximising the use of intensity information present in a spectrum. To this end we make use of fragmentation spectrum intensity predictions by MS2PIP^8^ to calculate an extensive set of features that provide additional discriminative and complementary information for PSM post-processing using machine learning. By comparing the intensity information in these predicted spectra with those recorded in the experimental spectra, our tool – ReScore – can be used to sensitively identify truthful PSMs while maintaining controlled FDR. This allows ReScore to go beyond explained peak counting or summing explained peak intensities: fragment ion peaks with predicted low intensities should have a positive contribution to the PSM score if that peak was indeed observed with low intensity, something that is not the case for current search engine PSM scoring functions.

We moreover show that these features are sufficient to render ReScore effectively independent of the original search engine score and related features, thus making ReScore compatible with any desired search engine, or with the combined output of multiple search engines, as for instance obtained by PeptideShaker^22^.

## Methods

### MS2PIP

MS2PIP is used to predict fragment ion intensities given a (modified) peptide sequence and charge. MS2PIP contains different models for different fragmentation types; definite models for singly charged fragment ions exist for two fragmentation types: HCD and CID. Models for EThcD fragmentation and for doubly charged fragment ions in HCD are also available in beta. Since its initial publication, MS2PIP has gone through several updates, and is currently available as a web-service^8^ and as a locally installable package^7^.

The input for MS2PIP is a PEPREC (PEPtide RECord^7^) file. This file contains peptide sequences, post-translational modifications (PTMs) and charges, each associated to a unique spectrum identifier. MS2PIP reads this file, and predicts the intensity for each theoretical fragment ion. If MS2PIP is provided with experimental spectra through an mgf file (mascot generic format^18^), and there is a correspondence between spectrum identifiers in the PEPREC file and the “TITLE” field of the spectrum file, it outputs both the experimental intensity (obtained from the mgf file) and the predicted intensity for each fragment ion.

### Feature Extraction & Selection

A machine learning method’s performance is strongly tied to the features used to describe the problem it will try to solve. Several metrics can be used to compare the experimental and empirical spectrum pairs that we have for each PSM, and a large number of features can therefore be devised.

MS2PIP predicted spectra contain *log2* transformed total-ion-current (TIC) normalized intensities for the theoretical fragment ions^8,9^. Given the nature of the *log2* transformation, low intensity peaks are weighed relatively more heavily by spectrum comparison metrics.

Spectral correlation features are therefore calculated twice: once on the *log2* transformed spectra, and once on the TIC-normalized spectra in the original linear scale. This allows different similarities and differences to be emphasized by the two distinct intensity scales. Moreover, these correlation calculations are not only performed for the entire spectrum but also for the b- and y-ion series separately. The resulting set of features is given in SI, T1. Note that this set of features will include significant redundancy. Despite the model used being an SVM, which includes regularization strategies that can handle issues that can stem from this redundancy, we choose to select features based on their correlations. By eliminating features that correlate to other features with a Pearson correlation coefficient of 0.9 or greater, we reduce the feature set to a total of 44 features (around 60% of its initial size). The selected features can be found in SI 5.

### Data sets & Processing

To validate the pipeline we propose, we present the result of an entrapment experiment. This type of experiment consists of adding protein sequences to the target database that correspond to real proteins that are not expected in the sample. Spectrum identification methods can be validated by comparing the FDR estimated by the decoy sequences with the fraction of entrapment (i.e., false) PSMs included in the results.

Our choice of entrapment is the established *Pyrococcus furiosus* (Pfu) standard for protein identification^21^, as obtained from the PRIDE archive^20,24^. A database of proteins was constructed by adding to the database with all Pfu sequences (both reviewed and unreviewed) all *Homo sapiens* sequences (reviewed), and all eukaryota sequences (reviewed). All of these were obtained via UniProt^6^. More details on the search settings can be found in SI 1 and SI 2.

The search results were converted into a Percolator input file (pin)^11^, which contains Percolator features for each PSM. These features can be split into two main groups: features based on the peptide sequence (such as amino-acid frequency, peptide length and precursor charge), and features based on the search engine’s score (such as, in this case, the scoring metrics calculated by MS-GF+^15^). Percolator is executed on these searches, so that a “baseline improvement” can be considered and compared against our proposed approach.

From the list of all PSMs obtained by MS-GF+, an input file for MS2PIP (that is, a PEPREC file) was created. This file was ran through MS2PIP, along with the mgf file containing the original spectra, producing a file with pairs of spectra (predicted and experimental) for each input PSM sequence. From these pairs, a matrix of features was calculated, where each row corresponds to a PSM and each column to a feature metric for that PSM from the table in SI 5. This matrix was combined with the pin file so that both the default Percolator features and the spectral comparison features are available for each PSM. Then Percolator was ran with this compound matrix as input. In addition, Percolator was also run using only the MS2PIP features, which thus notably excludes all MS-GF+ scoring information.

Besides these validation experiments, we show the results obtained when applying our procedure in two larger and more noisy datasets. These were obtained from PRIDE^5,14^; the first contains two a HEK cell line, and the second is a selection from the draft of the human proteome^13^ – specifically, all datasets from an adult adrenal gland. Search settings used were based on the information included in the PRIDE repository, and can be found in SI 3 and SI 4, respectively.

The code and data necessary to reproduce these results are available on GitHub^4^, along with installation and usage instructions.

## Results

### Entrapment experiments

The discriminative power of the spectral intensity prediction features computed by ReScore was evaluated by comparing the results of the MS-GF+ search engine with three different post-processing identification strategies, each using a different set of features: the default Percolator features, only MS2PIP-derived intensity correlation features, and the Percolator and MS2PIP-derived features combined. For each comparison we looked at the overlap in PSM identifications. To verify the method’s robustness, two entrapment experiments were performed as described in the previous section.

The first entrapment experiment consisted of the addition of all Homo sapiens sequences to the database the Pfu dataset is searched against. Figure 1 shows the outcome of these experiments, where each bar corresponds to one feature set.

**Figure 1:**
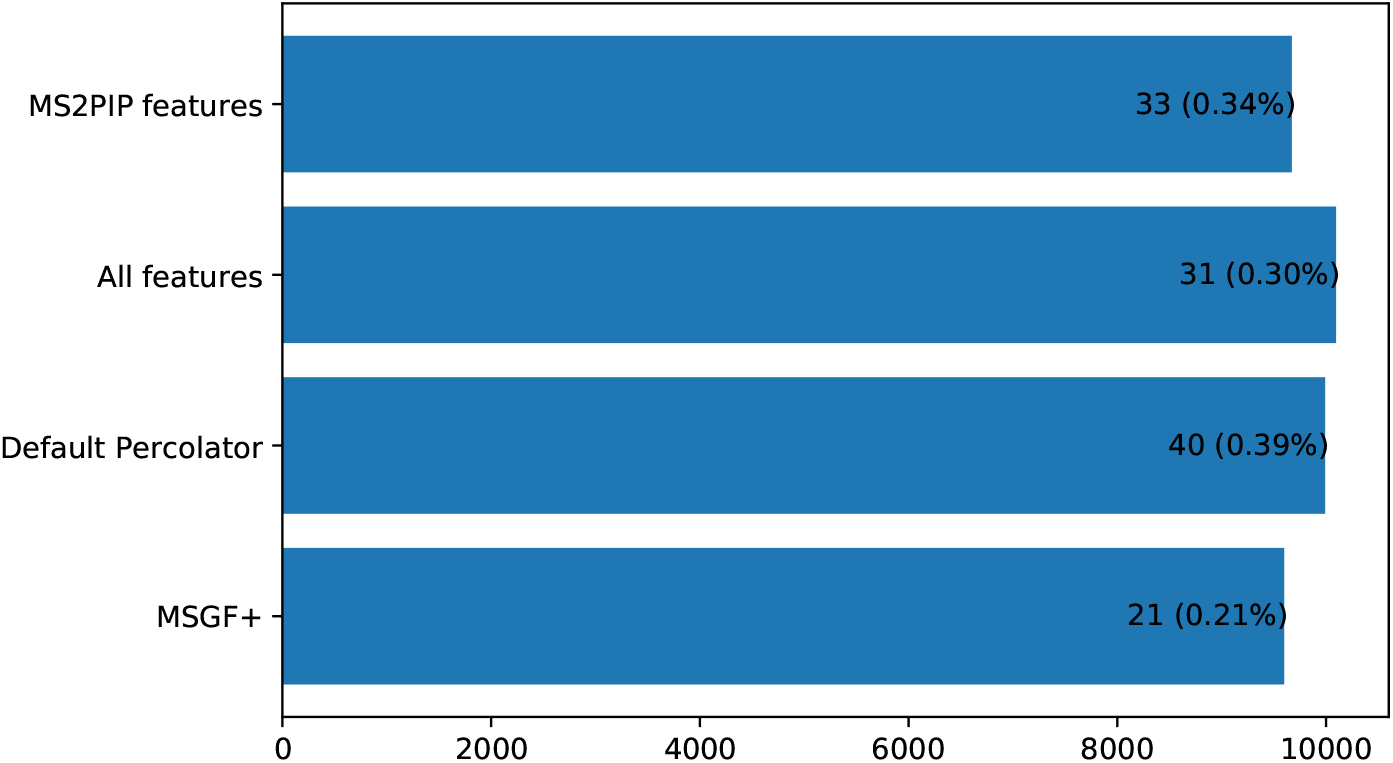
Number of identified PSMs (1% FDR) for each feature set (and for the search engine by itself), with amount and percentage of entrapment identifications written into each bar, for the *Homo sapiens* entrapment experiment.

With all four methods (that is, including the search engine), the amount of entrapment PSMs in the results is well under the FDR estimated with the decoy sequences; this indicates that no method over-fits to any particular characteristics of the decoy sequences, because if that were the case we would expect to see more entrapment sequences in the FDR-filtered results. Proceeding this experiment, a much larger entrapment database is used (as was also described previously). Large databases are known to give rise to issues when using classic spectrum identification techniques^16^, so a decrease in general performance is expected. However, the additional spectrum comparison-based features provide an additional layer of information that can be used to “rescue” identifications in these situations. The plot in Figure 2 shows the same comparison as Fig. 1 but for this second experiment.

**Figure 2:**
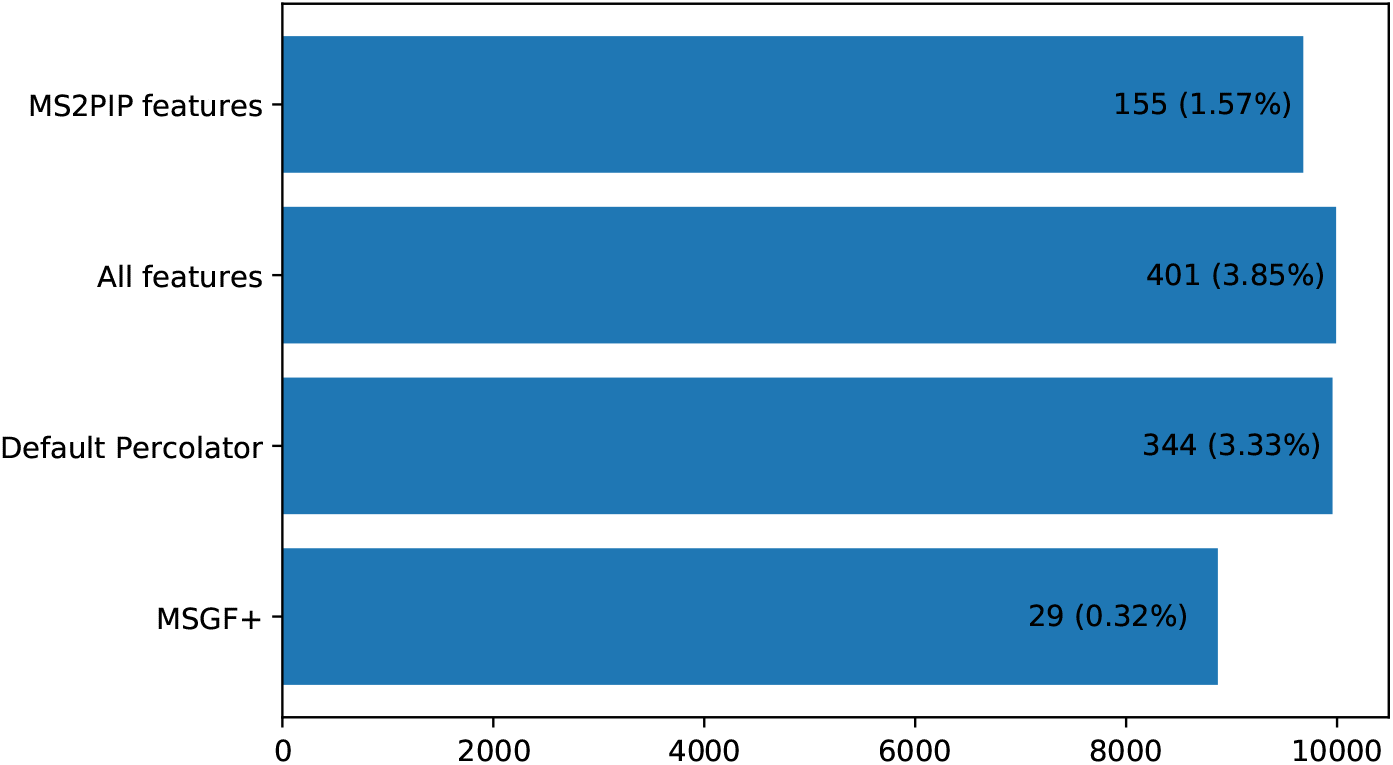
Number of identified PSMs (1% FDR) for each feature set (and for the search engine by itself), with amount and percentage of entrapment identifications written into each bar, for the eukaryota entrapment experiment.

Immediately it is noticed that the amount of PSMs obtained by MS-GF+ is much lower than previously; however, the search engine manages to control the FDR, maintaining the amount of entrapment PSMs well below the estimated 1%. While the three Percolator approaches manage to increase the number of PSMs to the range obtained in the previous experiment, they allow many more entrapment PSMs through, showing the effect of the database size increment in such approaches. However, using the MS2PIP-based features alone it is possible to obtain a balance, as we can observe an amount of PSMs on par as the previous experiment, and an amount of entrapment PSMs only marginally above the estimated FDR.

These two experiments show encouraging results. To further test and compare the performance of these three methods, results obtained from two other experiments are shown.

### Adult Adrenal Gland

The proposed framework was evaluated a larger dataset, downloaded from PRIDE under accession number PXD000561. As mentioned previously, one of the several tissues included in this experiment was selected and the spectra were searched as described in SI 4.

At 1% FDR, MS-GF+ reports the identification of 23988 PSMs. This number increases to 24995 when Percolator’s default feature set is used. Using the MS2PIP feature set, the number of PSMs reported at the same level of FDR is 24252. As seen previously, the largest increase in number of reported PSMs is seen when using the complete feature set; in that case, 25419 PSMs are reported at 1% FDR which translates an 6% increase. This analysis confirmed the previously observation that using the complete feature set is what returns more PSMs at a controlled FDR. The overlap between these three cases and the original search engine results can be seen in Fig. 3. To each feature set, three bars are associated: the orange bar represent the overlap between the rescored set of PSMs and the initial set obtained by MS-GF+; the other two bars represent the PSMs unique to each method (the search engine in blue, i.e., the “overruled” PSMs, and Percolator plus feature set in green – i. e., the “new” PSMs).

**Figure 3:**
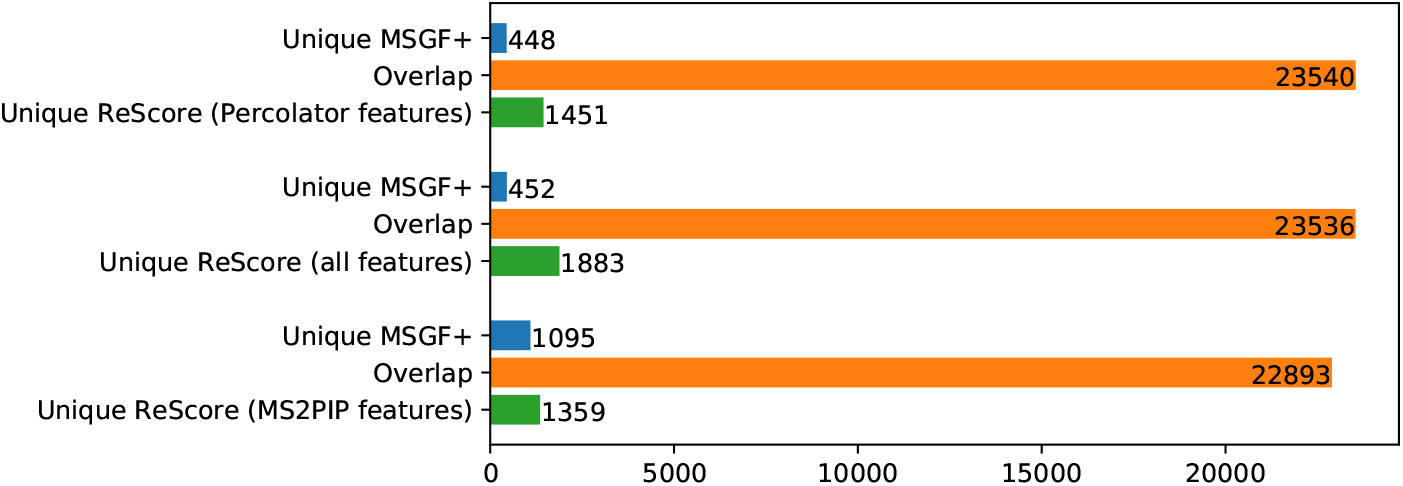
Number and overlap of identified PSMs (1% FDR) on the Adrenal Gland data set for each feature set, each compared with MS-GF+ as the baseline. Orange bars represent the overlap between post-processed PSM results and the MS-GF+ PSM results, green bars represent the amount of PSMs uniquely obtained by the post-processing method, while blue bars represent PSMs uniquely obtained by the search engine.

An additional analysis can be done concerning the “new” PSMs obtained with each method. In principle, one would expect that if the three methods pick up truthful PSMs, then these sets of “new” PSMs should show significant overlap. This is visualized in the Venn diagram in Figure 4

**Figure 4:**
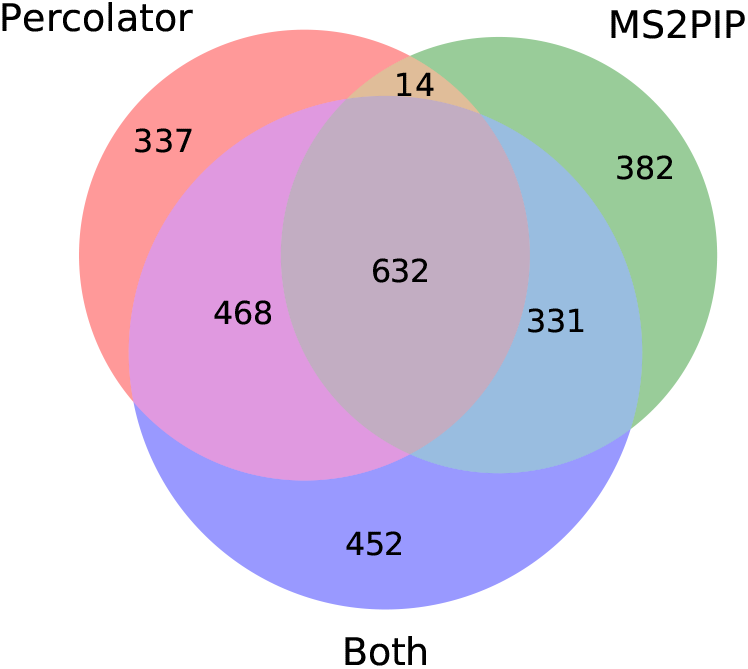
Relationship between the sets of PSMs obtained from the Adrenal Gland data set that each method adds to the results at 1% FDR.

Out of between 1359 and 1883 “new” PSMs, the three methods agree in 631 of them. Overall, in the case of the default Percolator feature set, about 77% of the added PSMs are supported by at least one other method; the same value is 71% for the MS2PIP feature set and 75% when using the combined feature set.

### HEK sample

For an assessment of ReScore performance on a much larger data set (1121149 spectra), the data for accession number PXD001468 was downloaded from the PRIDE database, which was processed as described in the methods section.

At 1% FDR, 467255 PSMs are reported by MS-GF+. Using Percolator’s default features set increased this number to 491833, thus adding a little more than 5%. When using only the MS2PIP features, the PSM number increases to 482096, which is an increase of 3%.

It should be noted that, as was the case for the two datasets discussed above, the highest number of PSMs (500270) is obtained when the default Percolator features are combined with the MS2PIP features. The overlap between the post processing results for the two feature sets and MS-GF+ alone are shown in Fig. 5, while the overlap in PSMs between the default Percolator feature set and the MS2PIP only feature set can be seen in Fig. 6.

**Figure 5:**
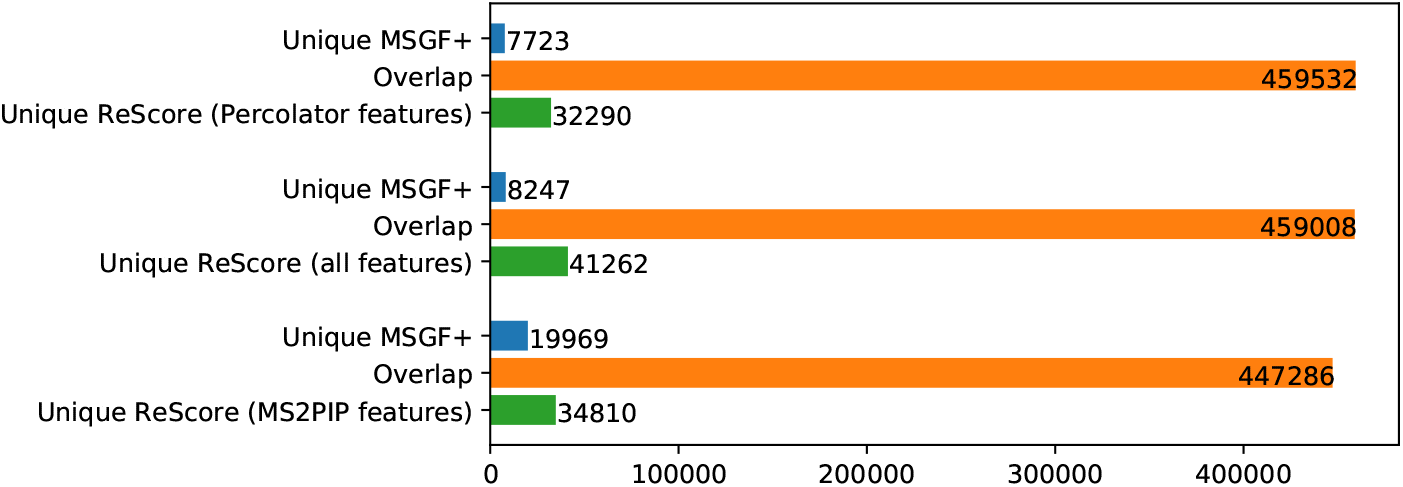
Number and overlap of identified PSMs (1% FDR) on the HEK data set for each feature set, each compared with MS-GF+ as the baseline. Orange bars represent the overlap between post-processed PSM results and the MS-GF+ PSM results, green bars represent the amount of PSMs uniquely obtained by the post-processing method, while blue bars represent PSMs uniquely obtained by the search engine.

**Figure 6:**
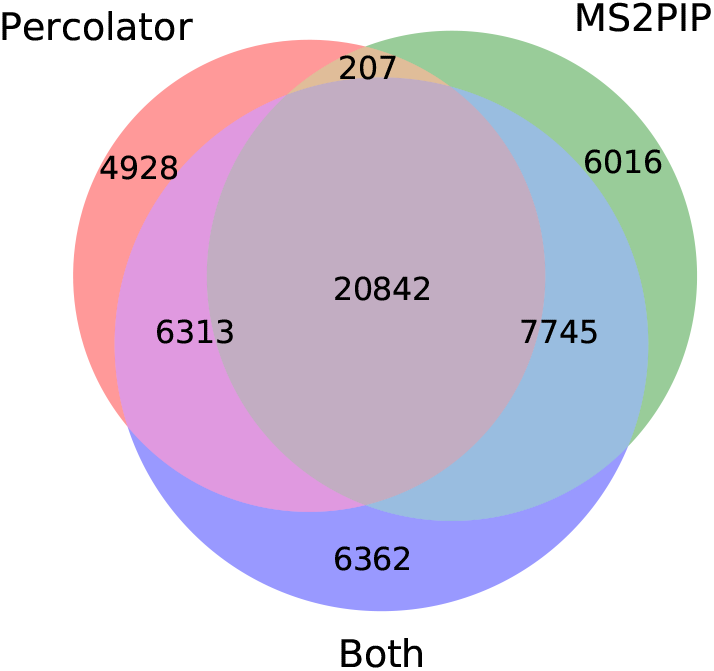
Relationship between the sets of PSMs obtained from the HEK data set that each method adds to the results at 1% FDR.

## Conclusion

In this work we have showed that, by comparing experimental spectra to theoretical spectra with fragment ion intensities computed by MS2PIP, Percolator can be applied independently from the search engine scoring function. Moreover, we showed that the resulting performance is on par with standard Percolator performance.

Therefore, we have showed that the post-processing step can effectively be decoupled from the search engine used to initially process the data: the only input this post-processing approach requires is a list of target and decoy PSMs, which can be obtained from any search engine or identification strategy.

Furthermore, we show consistently that the addition of our MS2PIP feature set to Percolator’s default feature set improves the amount of reported PSMs at a controlled level of FDR.

As a result, our study emphasizes how computationally predicted spectra can be used to replace static scoring functions with performant and adaptable machine learning algorithms.

