## Supporting Information for "Accurate peptide fragmentation predictions allow data driven approaches to replace and improve upon proteomics search engine scoring functions"

#### Table 1 - spectral comparison features

| | **Metric** | **Normalization** | **Full spectrum / b-ions / y-ions** | **Feature Name** |
|---|---|---|---|---|
| 1 | Pearson Correlation Coefficient | Normalized | full spectrum | spec_pearson |
| 2 | | | b-ions | ionb_pearson |
| 3 | | | y-ions | iony_pearson |
| 4 | | Logarithm of the normalized | full spectrum | spec_pearson_norm |
| 5 | | | b-ions | ionb_pearson_norm |
| 6 | | | y-ions | iony_pearson_norm |
| 7 | Spearman Correlation Coefficient | Normalized | full spectrum | spec_spearman |
| 8 | | | b-ions | ionb_spearman |
| 9 | | | y-ions | iony_spearman |
| 13 | Cosine | Normalized | full spectrum | cos |
| 14 | | | b-ions | cos_ionb |
| 15 | | | y-ions | cos_iony |
| 1 | | Logarithm of | full spectrum | cos_norm |

| | | | | |
|---|---|---|---|---|
| 6 | | the normalized | | |
| 17 | | | b-ions | cos_ionb_norm |
| 18 | | | y-ions | cos_iony_norm |
| 19 | Dot product | Normalized | full spectrum | dotprod |
| 20 | | | b-ions | dotprod_ionb |
| 21 | | | y-ions | dotprod_iony |
| 22 | | Logarithm of the normalized | full spectrum | dotprod_norm |
| 23 | | | b-ions | dotprod_ionb_norm |
| 24 | | | y-ions | dotprod_iony_norm |
| 25 | Mean Squared Error | Normalized | full spectrum | spec_mse |
| 26 | | | b-ions | ionb_mse |
| 27 | | | y-ions | iony_mse |
| 28 | | Logarithm of the normalized | full spectrum | spec_mse_norm |
| 29 | | | b-ions | ionb_mse_norm |
| 30 | | | y-ions | iony_mse_norm |
| 31 | Distribution of differences between predicted and empirical intensities | Normalized | full spectrum | min_abs_diff |
| 32 | | | | max_abs_diff |
| 33 | | | | mean_abs_diff |
| 34 | | | | std_abs_diff |
| 37 | | | b-ions | ionb_min_abs_diff |

| | | | | |
|---|---|---|---|---|
| 38 | | | | ionb_max_abs_diff |
| 39 | | | | ionb_mean_abs_diff |
| 40 | | | | ionb_std_abs_diff |
| 41 | | | y-ions | iony_min_abs_diff |
| 42 | | | | iony_max_abs_diff |
| 43 | | | | iony_mean_abs_diff |
| 44 | | | | iony_std_abs_diff |
| 45 | | Logarithm of the normalized | full spectrum | min_abs_diff_norm |
| 46 | | | | max_abs_diff_norm |
| 47 | | | | mean_abs_diff_norm |
| 48 | | | | std_abs_diff_norm |
| 49 | | | | min_abs_diff_iontype_norm |
| 50 | | | | max_abs_diff_iontype_norm |
| 51 | | | b-ions | ionb_min_abs_diff_norm |
| 52 | | | | ionb_max_abs_diff_norm |
| 53 | | | | ionb_mean_abs_diff_norm |
| 54 | | | | ionb_std_abs_diff_norm |
| 55 | | | y-ions | iony_min_abs_diff_norm |
| 56 | | | | iony_max_abs_diff_norm |
| 57 | | | | iony_mean_abs_diff_norm |

| | | | | |
|---|---|---|---|---|
| 58 | | | | iony_std_abs_diff_norm |
| 59 | | Normalized | full spectrum | abs_diff_Q1 |
| 60 | | | | abs_diff_Q2 |
| 61 | | | | abs_diff_Q3 |
| 62 | | Logarithm of the normalized | | abs_diff_Q1_norm |
| 63 | | | | abs_diff_Q2_norm |
| 64 | | | | abs_diff_Q3_norm |
| 65 | | Normalized | b-ions | ionb_abs_diff_Q1 |
| 66 | | | | ionb_abs_diff_Q2 |
| 67 | | | | ionb_abs_diff_Q3 |
| 68 | | Logarithm of the normalized | | ionb_abs_diff_Q1_norm |
| 69 | | | | ionb_abs_diff_Q2_norm |
| 70 | | | | ionb_abs_diff_Q3_norm |
| 71 | | Normalized | y-ions | iony_abs_diff_Q1 |
| 72 | | | | iony_abs_diff_Q2 |
| 73 | | | | iony_abs_diff_Q3 |
| 74 | | Logarithm of the normalized | | iony_abs_diff_Q1_norm |
| 75 | | | | iony_abs_diff_Q2_norm |
| 76 | | | | iony_abs_diff_Q3_norm |

#### 1. *Pyrococcus furiosus* + *Homo sapiens* entrapment search details

MS-GF+ v2017.01.13

Databases obtained from uniprot on June 20th 2018: reviewed and unreviewed sequences, concatenated with all reviewed *Homo sapiens* sequences

MS-GF+ search command:
java -Xmx28000M -jar MSGFPlus.jar modification.txt -s Velos05137.mgf -d PyrFu_human.fasta -o Velos05137.mzid -t 10ppm -tda 1 -m 3 -inst 1 -minLength 8 -minCharge 2 -maxCharge 4 -n 1 -addFeatures 1 -protocol 0 -thread 23

modification.txt: Carbamidomethyl and oxidation, maximum 2

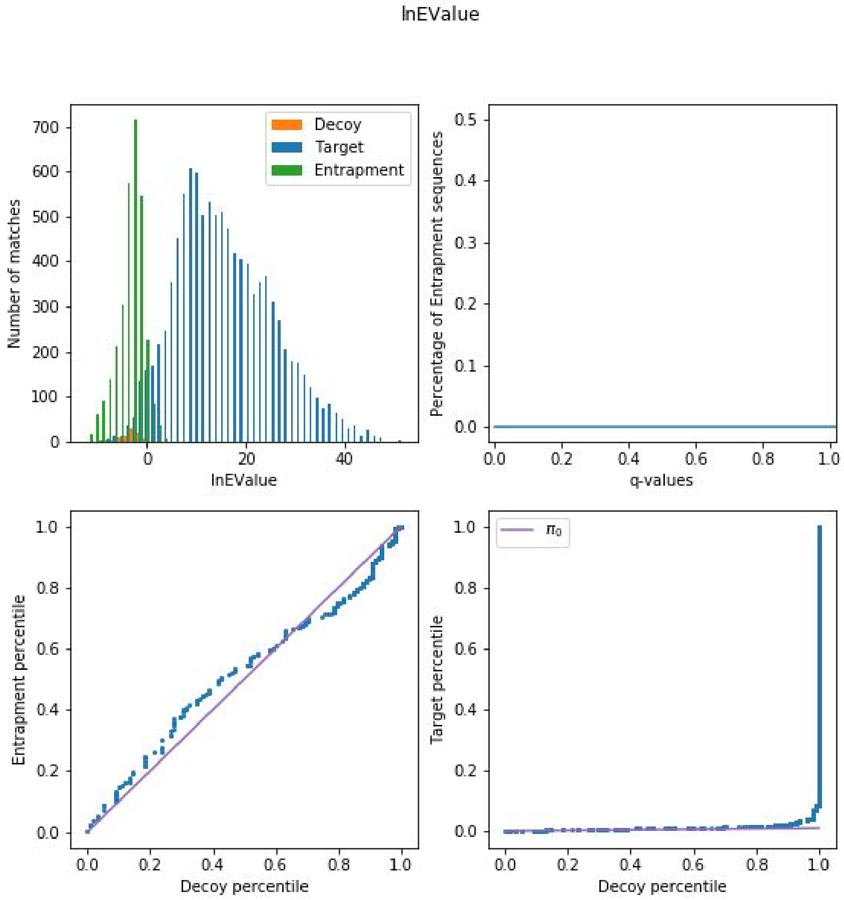

### 2. *Pyrococcus furiosus* + eukaryota entrapment search details

MS-GF+ v2017.01.13

Databases obtained from uniprot on June 20th 2018: reviewed and unreviewed sequences, concatenated with all reviewed eukaryota sequences

MS-GF+ search command:
java -Xmx28000M -jar MSGFPlus.jar modification.txt -s Velos05137.mgf -d PyrFu_euk.fasta -o Velos05137.mzid -t 10ppm -tda 1 -m 3 -inst 1 -minLength 8 -minCharge 2 -maxCharge 4 -n 1 -addFeatures 1 -protocol 0 -thread 23

modification.txt: Carbamidomethyl and oxidation, maximum 2

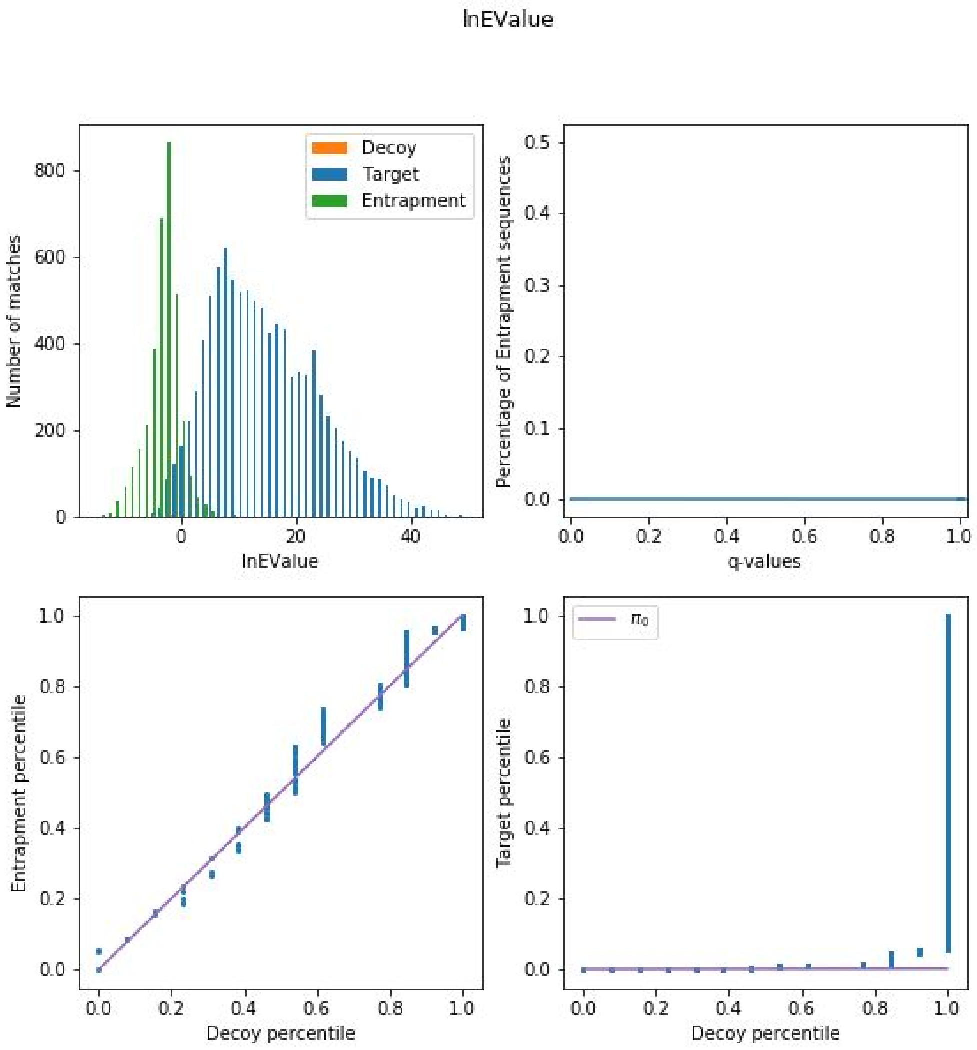

### 3. HEK sample (PXD001468) search details

https://www.ebi.ac.uk/pride/archive/projects/PXD001468
All experiments concatenated into one large mgf file, which was posteriorly searched. The concatenated file is available here.

MS-GF+ v2017.01.13

Database of human proteins, MQ contaminants and crap database

MS-GF+ search command:
java -Xmx90000M -jar MSGFPlus.jar modification.txt -s all.mgf -d database.fasta -o all.mzid -t 10ppm -tda 1 -m 3 -inst 1 -minLength 8 -minCharge 2 -maxCharge 4 -n 1 -addFeatures 1 -protocol 0 -thread 23

modification.txt: Carbamidomethyl and oxidation, maximum 2

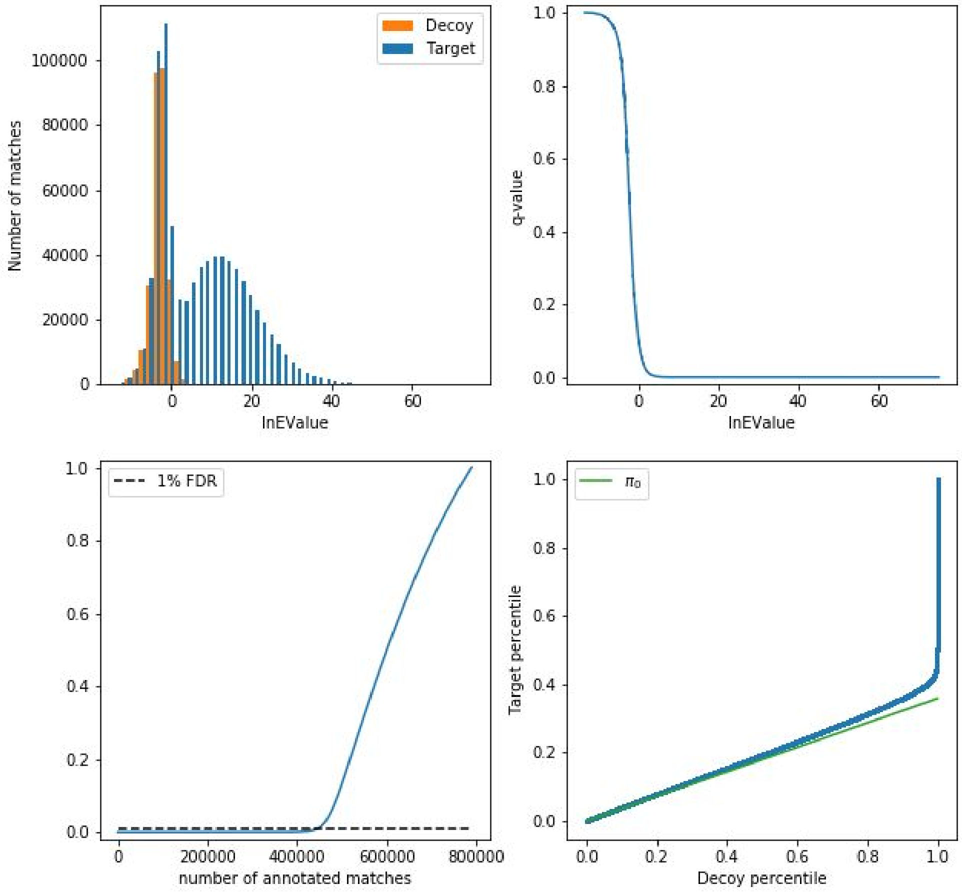

#### 4. Adult Adrenal Gland (PXD000561) search details

https://www.ebi.ac.uk/pride/archive/projects/PXD000561
All adrenal gland experiments concatenated into one large mgf file, which was posteriorly searched. The concatenated file is available here.

MS-GF+ v2017.01.13

Database of human proteins, MQ contaminants and crap database

MS-GF+ search command:
java -Xmx90000M -jar MSGFPlus.jar modification.txt -s all.mgf -d synth.fasta -o all.mzid -t 10ppm -tda 1 -m 3 -inst 1 -minLength 8 -minCharge 2 -maxCharge 4 -n 1 -addFeatures 1 -protocol 0 -thread 23

modification.txt: Oxidation and phosphorylation, maximum 2

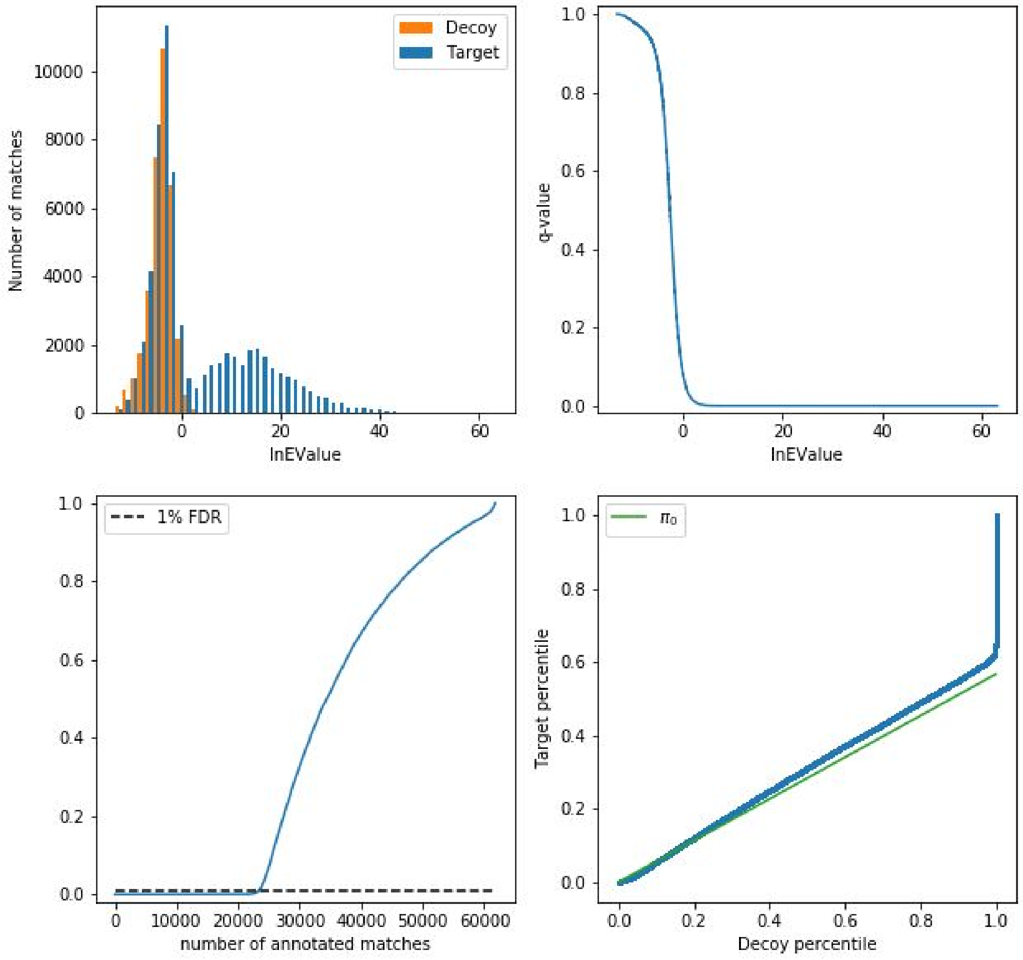

### 5. Selected MS2PIP features

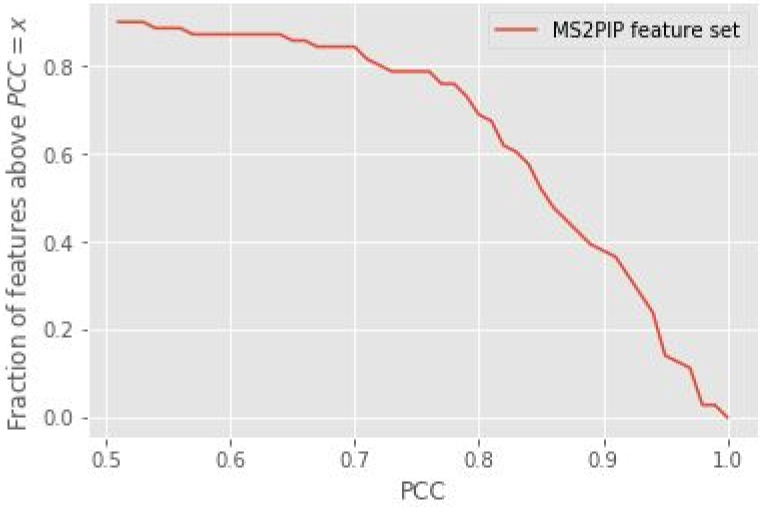


- iony_pearson_norm,
- spec_pearson,
- spec_pearson_norm,
- iony_pearson,
- ionb_pearson_norm,
- dotprod_ionb,
- dotprod_norm,
- dotprod,
- cos_norm,
- cos_iony,
- cos_iony_norm,
- cos_ionb_norm,
- ionb_mse,
- spec_mse_norm,
- spec_mse,
- ionb_mse_norm,
- min_abs_diff,
- min_abs_diff_norm,
- min_abs_diff_iontype,
- iony_min_abs_diff,
- ionb_mean_abs_diff,
- mean_abs_diff,
- max_abs_diff,
- max_abs_diff_norm,
- max_abs_diff_iontype,
- abs_diff_Q1,

- abs_diff_Q1_norm,
- abs_diff_Q2,
- abs_diff_Q2_norm,
- abs_diff_Q3,
- ionb_max_abs_diff,
- ionb_max_abs_diff_norm,
- ionb_min_abs_diff,
- ionb_min_abs_diff_norm,
- ionb_abs_diff_Q1,
- ionb_abs_diff_Q1_norm,
- ionb_abs_diff_Q2,
- ionb_abs_diff_Q2_norm,
- ionb_abs_diff_Q3,
- ionb_abs_diff_Q3_norm,
- iony_abs_diff_Q1,
- iony_abs_diff_Q2,
- iony_abs_diff_Q3,
- iony_min_abs_diff_norm.
